# Disruption of consciousness depends on insight in OCD and on positive symptoms in schizophrenia

**DOI:** 10.1101/2024.01.02.571832

**Authors:** Selim Tumkaya, Bengü Yücens, Muhammet Gündüz, Maxime Maheu, Lucie Berkovitch

## Abstract

Disruption of conscious access contributes to the advent of psychotic symptoms in schizophrenia but could also explain lack of insight in other psychiatric disorders.

In this study, we explored how insight and psychotic symptoms related to disruption of consciousness. We explored consciousness in patients with schizophrenia, patients with obsessive-compulsive disorder (OCD) with good vs. poor insight and matched controls. Participants underwent clinical assessments and performed a visual masking task allowing us to measure individual consciousness threshold. We used a principal component analysis to reduce symptom dimensionality and explored how consciousness measures related to symptomatology.

We found that clinical dimensions could be well summarized by a restricted set of principal components which also correlated with the extent of consciousness disruption. More specifically, positive symptoms were associated with impaired conscious access in patients with schizophrenia whereas the level of insight delineated two subtypes of OCD patients, those with poor insight who had consciousness impairments similar to patients with schizophrenia, and those with good insight who resemble healthy controls.

Our study provides new insights about consciousness disruption in psychiatric disorders, showing that it relates to positive symptoms in schizophrenia and with insight in OCD. In OCD, it revealed a distinct subgroup sharing neuropathological features with schizophrenia. Our findings refine the mapping between symptoms and cognition, paving the way for a better treatment selection.

## INTRODUCTION

Consciousness can be defined as the ability to generate and maintain mental representations.^1^ Across various experimental paradigms, patients with schizophrenia were shown to have an impaired conscious access.^2^ They need, for instance, a longer delay to consciously perceive an external stimulus compared to healthy participants.^3^ Interestingly, a disruption of conscious access may foster psychotic symptoms in schizophrenia, as well as in other nosographical categories, such as bipolar disorder.^4^ Neurological disorders devoid of psychotic symptoms also exhibit elevated consciousness threshold,^5–7^ suggesting that conscious access impairments could translate into various other symptoms such as cognitive impairments and anosognosia.

To explore this hypothesis, we investigated conscious access in patients with obsessive-compulsive disorder (OCD). Indeed, patients with OCD have variable levels of insight, and OCD with poor insight has been related to specific cognitive dysfunctions, some of which being shared with schizophrenia.^8^ Clinical scales, such as the Over Valued Ideas Scale (OVIS) can be used to measure insight in patients with OCD.^8–13^ Notably, OVIS measures overvalued ideas, including items measuring to which extent beliefs are “unshakable.”

In the current study, we intended to explore how conscious access and processing vary according to psychotic symptoms in patients with schizophrenia, on the one hand, and with the level of insight and belief strength in patients with OCD on the other hand. We expected that patients with OCD having high OVIS scores (i.e., strong beliefs and poor insight) would have an impaired conscious access close to that of patients with schizophrenia, whereas conscious access in patients with low OVIS scores (weak beliefs and good insight) would resemble that of healthy controls. We recruited patients with OCD, with various levels of insight, patients with schizophrenia and healthy controls, who all performed a visual masking task which allows quantification of conscious and non-conscious processing in controlled settings. In addition, patients were submitted to relevant clinical scales (e.g., OVIS for OCD and SAPS for schizophrenia) and we used a principal component analysis (PCA) to reduce symptom dimensionality and explore how symptomatology axes related to consciousness measures as derived from the visual masking task.

We found that meaningful clinical dimensions were identified by PCA-decomposition of clinical scales, which also predicted impairments of consciousness in patients. More specifically, positive symptoms were associated with impaired conscious access in patients with schizophrenia whereas the level of insight delineated two different subtypes of patients with OCD, those with poor insight who had disruption of consciousness similar to patients with schizophrenia and those with good insight who were comparable to healthy controls.

## MATERIAL AND METHODS

### Participants

We recruited 60 people: 20 patients with OCD, 20 patients with psychosis (schizophrenia or schizoaffective disorder) according to DSM-5 criteria from Pamukkale University Faculty of Medicine, Psychiatry Hospital Polyclinics, and 20 healthy volunteers who were free from any psychiatric or neurological disease and matched to patients in terms of age, gender and education (see Table 1). All participants give their informed consent after receiving a complete description of the study. The study was approved by Pamukkale University Clinical Research Ethics Committee (decision n°60116787-020/49020).

**Table 1.** Participants’ characteristics.

| Characteristic | Controls,<br>N = 19<sup>a</sup> | OCD with good<br>insight,<br>N = 10<sup>a</sup> | OCD with poor<br>insight,<br>N = 9<sup>a</sup> | Schizophrenia,<br>N = 17<sup>a</sup> | P-value<sup>b</sup> |
| --- | --- | --- | --- | --- | --- |
| **Gender (F/M)** | 11/8 | 5/5 | 7/2 | 8/9 | 0.5 |
| **Age (years)** | 34.05 (1.86) | 28.80 (2.82) | 35.78 (3.17) | 38.00 (2.69) | 0.12 |
| **Smoking** | 7 (37%) | 1 (10%) | 0 (0%) | 5 (29%) | 0.11 |
| **Education (years)** | 14.00 (0.56) | 14.80 (0.94) | 12.22 (0.97) | 12.82 (0.70) | 0.14 |
| **Number of**<br>**hospitalization(s)** | NA | 0.00 (0.00) | 0.44 (0.34) | 3.35 (0.84) | 0.002 |
| **Duration of the disease**<br>**(years)** | NA | 12.10 (2.76) | 16.00 (2.96) | 14.12 (2.58) | 0.7 |
| **Fluoxetine equivalent**<br>**(mg/day)** | NA | 25.29 (4.99) | 45.08 (9.12) | 12.73 (6.67) | 0.013<sup>c</sup> |
| **Olanzapine equivalent**<br>**(mg/day)** | NA | 0.00 (0.00) | 0.00 (0.00) | 23.30 (3.52) | <0.001<sup>c</sup> |
| **YBOCS (total score)** | NA | 17.50 (2.45) | 22.56 (1.83) | NA | 0.12 |
| **DOCS (total score)** | NA | 22.20 (4.27) | 29.22 (4.13) | NA | 0.3 |
| **HAM-D (total score)** | NA | 4.10 (0.86) | 9.00 (3.02) | NA | 0.12 |
| **HAM-A (total score)** | NA | 7.50 (1.73) | 9.56 (3.60) | NA | 0.6 |
| **OVIS (total score)** | NA | 4.18 (0.27) | 6.47 (0.13) | NA | <0.001 |
| **SAPS (total score)** | NA | NA | NA | 9.00 (2.58) | NA |
| **SANS (total score)** | NA | NA | NA | 29.71 (5.03) | NA |
| **Calgary Depression Scale**<br>**(total score)** | NA | NA | NA | 3.29 (0.62) | NA |

<sup>a</sup>n (%); Mean (sem)
<sup>b</sup>Pearson's Chi-squared test; One-way ANOVA
<sup>c</sup>P-values calculated only between the three groups of patients
Y-BOCS: Yale-Brown Obsession Compulsion Scale (Y-BOCS); DOCS: Dimensional Obsession Compulsion Scale; HAM-D: Hamilton Depression Rating Scale; HAM-A: Hamilton Anxiety Rating Scale; OVIS: Over Valued Ideas Scale (OVIS); SAPS: Scale for the Assessment for Negative Symptoms; SANS: Scale for the Assessment of Negative Symptoms.

### Procedure

#### Clinical assessments

In the OCD group, we used the Yale-Brown Obsession Compulsion Scale (Y-BOCS),^14^ the Dimensional Obsession Compulsion Scale (DOCS),^15^ the Hamilton Depression Rating Scale (HAM-D),^16^ the Hamilton Anxiety Rating Scale (HAM-A),^17^ and the Over Valued Ideas Scale (OVIS),^9^ which was used to split the OCD group into two subgroups: OCD with weak beliefs or good insight (OVIS < 6) and OCD with strong beliefs or poor insight (OVIS ≥ 6).^8–13^ In the psychosis group, we used the Scale for the Assessment for Negative Symptoms (SAPS), the Scale for the Assessment of Negative Symptoms (SANS)^18^ and the Calgary Depression Scale (CDS).^19^ Doses of prescribed antipsychotic and antidepressant treatments were converted to olanzapine^20^ and fluoxetine equivalents.^21^

#### Visual backward masking paradigm

After the clinical evaluation, participants performed a visual masking task. Visual masking experiments consist in displaying a short-duration visual stimulus that is subsequently masked by a second stimulus. As the delay between the two stimuli (stimulus onset asynchrony, SOA) increases, the first stimulus becomes easier to perceive. In the current experiment, we adapted a visual sequence, used in previous studies,^3,4,22^ where a target digit (2, 3, 7, or 8) is masked by a group of letters.

Target digit visibility was parametrically manipulated with eight target-mask delays (16.7, 33.3, 50, 66.7, 83.3, 100, 116.7 or 166.7 ms) that were randomly intermixed across trials (**Fig. 1**). In 20% of the trials, the target digit was replaced by a blank with the same duration (catch trials), so we could measure participants’ ability to detect the presence of the digit. Each participant performed a training block of 20 trials, followed by 2 blocks of 160 trials each, separated by a break. Our experiment was coded with *MATLAB* and *Psychtoolbox.*^23,24^

**Fig. 1.**
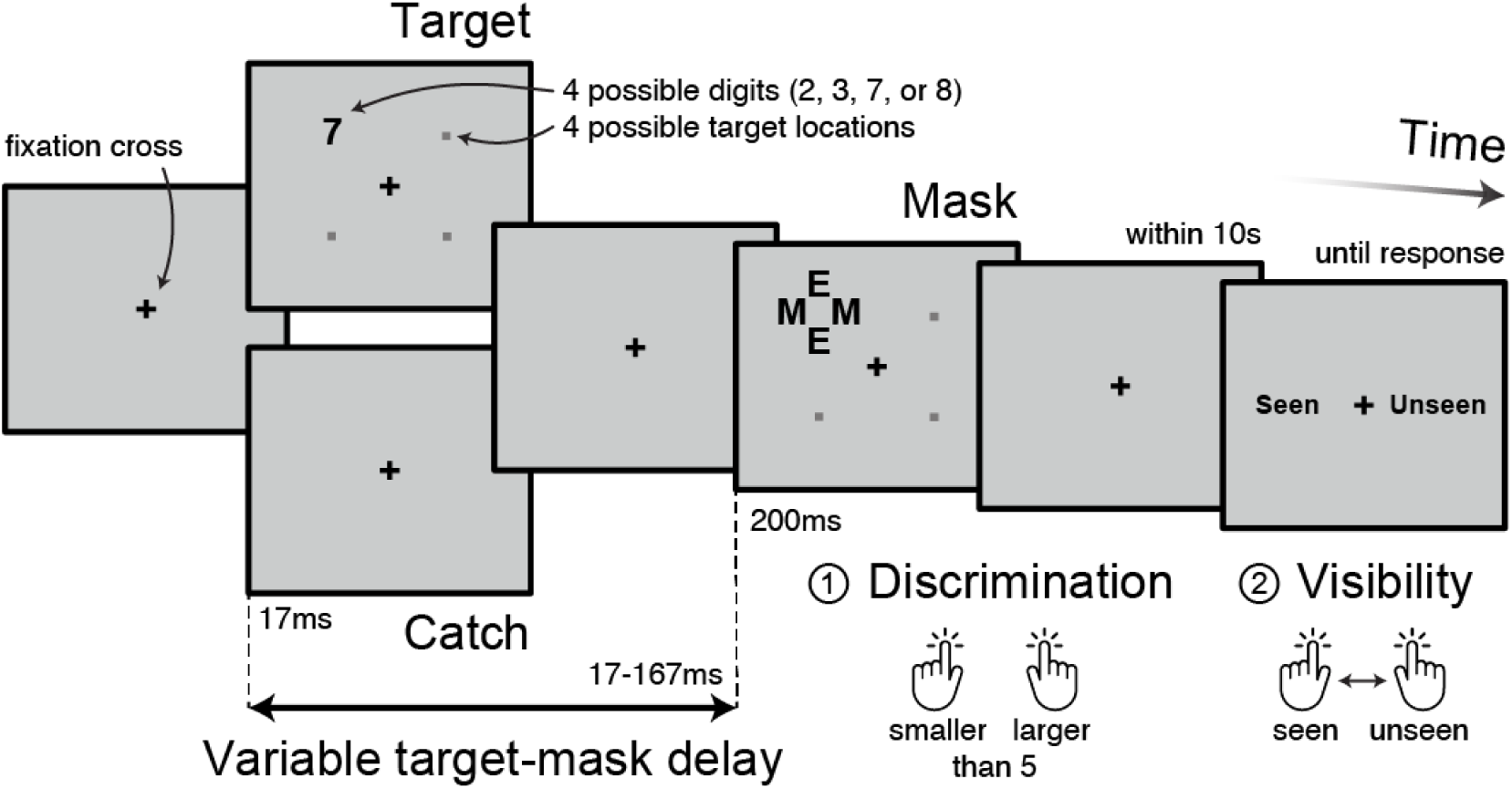
Visual backward masking paradigm. A fixation cross was displayed in the center of the screen during approximately one second (randomly jittered across trials). A target digit (2, 3, 7, or 8) was then presented for a fixed duration of 16.7 ms at a random position among four (1.4 degrees above or below and 1.4 degrees right or left of the fixation cross). In 20% of the trials, the target digit was replaced by a blank of the same duration (catch trials). After a variable delay (stimulus onset asynchrony, SOA), a metacontrast mask appeared at the target location for 200 ms. The mask was composed of four letters (two horizontally aligned M and two vertically aligned E) surrounding the target stimulus location without superimposing or touching it. Target digit visibility was parametrically manipulated by using eight possible target-mask delays (16.7, 33.3, 50, 66.7, 83.3, 100, 116.7 or 166.7 ms) that were randomly intermixed across trials. Participants had at most 10 seconds to determine whether the number was smaller or greater than 5, by pressing “S” or “L” respectively on the keyboard. Then, they had to rate the subjective visibility of the digit. The response words “seen” and “unseen” randomly appeared on the right and the left sides of the fixation cross and participants responded by pressing the button (“S” or “L”) corresponding to the side of the response they wanted to select. The two alternatives remained on screen until a response was made.

This paradigm with a high sampling of target-mask delays was designed as an alternative to staircase procedures, which adjust target-mask delays to participant’s accuracy (in detecting and/or discriminating the target) thus converging on individual consciousness threshold.^4,5,25,26^ Our whole-range, parametric screening of different target-mask delays provides useful additional information over staircase procedures. In particular, it allows probing the integrity of conscious and unconscious processing as well as the relationships between discrimination and detection abilities.

### Cognitive and clinical variables

#### Consciousness measures

For each participant, we measured the accuracy in comparing the target against 5 (discrimination) and the rate of seen trials among non-catch trials (visibility). An objective measure of conscious access was computed by confronting subjective visibility (seen versus not seen) against the presence or absence of a target (target versus catch trials), resulting in a *d*-prime measure (detection). Conscious trials were defined as trials where digits were both rated as “seen” and correctly categorized. All these measures of conscious access usually increase non-linearly with target-mask delay (stimulus onset asynchrony, SOA), following a sigmoid curve^3,27^ which we fitted using the logistic model:

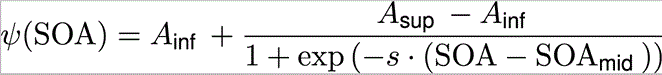

where *A*_inf_ is the lower asymptote, *A*_sup_ the upper asymptote, SOA_mid_ the inflection point and *s* the slope. Parameter values were determined with gradient-based optimization. Whenever the fit did not converge, starting values were manually set after visual eyeballing of the participant’s data. The quality of the fit was estimated by the proportion of variance explained (*R*²).

In line with staircase paradigm studies,^4,5^ the consciousness threshold was estimated as value of target-mask delay corresponding to 50% of “seen and correct” trials in the model, that is *ψ*(SOA) = 0.5. Similarly, visibility threshold was defined by *ψ*(SOA) = 0.5 based on “seen” trials, and discrimination threshold by *ψ*(SOA) = 0.75 for “correct” trials (since chance level is at 0.5 for discrimination).

Previous studies showed that participants with schizophrenia have alterations of both conscious access (elevated threshold to consciously perceive information) and conscious processing (lower accuracy than controls even for long target-mask delays, when information is supposed to be consciously perceived), whereas unconscious processing is preserved.^2,3,22^ In our model, the integrity of conscious processing can be assessed by the upper asymptote (*A*_sup_) and the integrity of unconscious processing by the lower asymptote (*A*_inf_). While altered conscious processing has already been observed in schizophrenia^3,22^ and could reflect attentional lapses^28^ or noisier processing,^29,30^ differences in upper asymptotes could drive potential differences in consciousness threshold. In particular, a decrease of the upper asymptote could lead to an underestimation of the consciousness threshold. We therefore run the analyses before and after scaling each participant’s data according to their upper asymptote resulting in a proportion of conscious trials equals to 1 at the longer target-mask delays. This allowed to compute a corrected consciousness threshold, independent from alterations in conscious processing or attentional lapses.^28^

#### Additional measures

Patients with schizophrenia and patients with OCD generally exhibit alterations of metacognitive processing, notably in perceptual decision-making tasks.^31–34^ To assess metacognitive abilities in our task, we measured the difference in discrimination accuracy between seen and unseen trials (which should ideally equal 100% for seen and 50% for unseen trials).

Even if participants were not instructed to answer as fast as possible, we analyzed reaction times to ensure that any observed variations in consciousness measures would not relate to differences in speed/accuracy trade-off.^35^

#### Principal component analysis of symptom dimensions

To assess the clinical profile of our sample of patients, we run a PCA on all items from all clinical scales, separately for the schizophrenia and OCD group, using the same approach and analytical tools as previous studies.^36,37^ Principal components (PC) capture how the symptoms are correlated (or anticorrelated) one with another, thereby reflecting entirely data-driven symptomatic dimensions within a group. Each item was first scaled to have unit variance across patients, and items having null variance were discarded. Significance of PC was computed via permutation-based non-parametric testing. At each permutation, patient order was randomly shuffled for each symptom variable before the PCA was re-computed. Permutations were repeated 10,000 times to establish the null distribution. PCs which accounted for a proportion of variance that exceeded chance (*P <* 0.05 across all permutations) were retained for further analysis, and ordered as a function of the proportion of symptom variance they accounted for.

### Statistical analyses

Analyses of variance (ANOVAs) were conducted after excluding extreme reaction times (below 200 ms and above 6 seconds) on each consciousness measure, with target-mask delays as a within-subject factor and group as a between-subject factor. ANOVAs were performed across participants, to examine the effect of age, gender and smoking on behavioral measures of conscious access. ANOVAs and Pearson correlations were conducted at the group level to measure the interactions between measures of conscious access (consciousness threshold, upper asymptotes, discrimination and visibility thresholds, discrimination accuracy, detection *d*-primes, seen vs. unseen difference in discrimination accuracy) and several variables including clinical measures (all above-mentioned clinical scales, PC scores, duration of disease), treatment posology (olanzapine and fluoxetine equivalents), with covariates (target-mask delays, age, gender and smoking status) whenever they appear to significantly interact with consciousness measures across participants, or using a simple correlation otherwise. Given their central role in our working hypothesis, the significance of correlation coefficients between key clinical scales (OVIS and SAPS) or PC scores, on the one hand, and consciousness threshold, on the other hand, was confirmed by permutation-based non-parametric testing (10,000 permutations; *P <* 0.05).

All statistical analyses were two-tailed and performed using the *R* statistical software (https://www.r-project.org). Whenever we wanted to explore the absence of differences, we run Bayesian statistics using the *BayesFactor* package.^38^ We directly reported the inverse Bayes factor (BF) quantifying the evidence in favor of the null compared to the alternative hypothesis (e.g., when 1/BF = 4.0, it means that the null is four times more plausible than the alternative hypothesis).

### Exclusion of participants

Five participants (three with schizophrenia, one with OCD poor insight and one healthy control) had to be excluded from all the consciousness measures analyses because they had abnormally low or high performances for both discrimination and visibility. Four additional participants (two healthy controls, one patient with OCD and good insight, one patient with OCD and poor insight) did not correctly perform one of the two tasks (three were at chance-level for the discrimination task and one reported all trials as “seen”), whereas their results for the other task were comparable to that of the other participants, suggesting that they misunderstood instructions. We excluded the data of the task that they failed to perform.

## RESULTS

### Parametric modulation of conscious perception by visual masking

In our paradigm, measures of consciousness are expected to parametrically increase with target-mask delays and this is indeed what we observed for discrimination accuracy (*F*_7,336_ = 361.91, *P <* 0.001), visibility (*F*_7,350_ = 580.07, *P <* 0.001), and detection *d*-prime (*F*_7,350_ = 190.15, *P <* 0.001).

### Impaired discrimination and visibility in schizophrenia and OCD with poor insight

We aimed to replicate past findings demonstrating a disruption of consciousness in patients with schizophrenia. Accordingly, we observed that schizophrenia patients had lower discrimination accuracy (*F*_1,32_ = 14.75, *P <* 0.001) and visibility (*F*_1,34_ = 11.88, *P =* 0.002) than healthy controls (see **Fig. 2A**). In line with our main working hypothesis, we then observed that patients with OCD exhibit consciousness impairment which depended on their level of insight: OCD patients with poor insight (OVIS ≥ 6) had a lower discrimination accuracy than healthy controls (*F*_1,24_ = 11.51, *P =* 0.002), whereas OCD patients with good insight (OVIS < 6) had a discrimination accuracy similar to that of healthy controls (*F*_1,24_ = 0.27, *P =* 0.61, 1/BF = 6.1). This result was also observed for visibility and detection *d*-prime (**Supplementary Table 1**). Crucially, there was no significant difference in all consciousness-related measures considered between patients with schizophrenia and patients with OCD and poor insight (all 1/BF > 4).

**Fig. 2.**
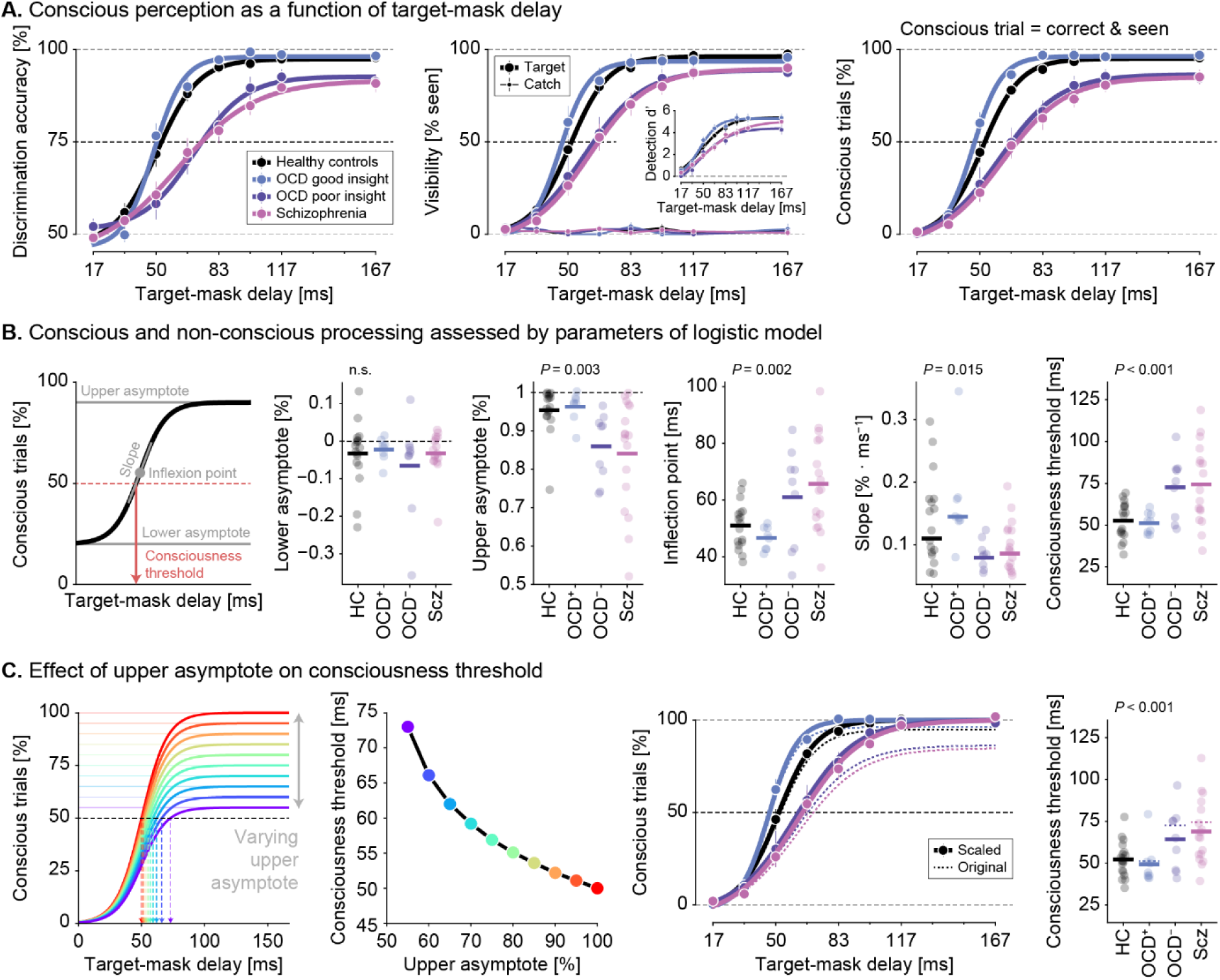
Impairments of conscious perception across groups. **A**. Left, discrimination accuracy (i.e., comparing the masked digit to 5); Middle, visibility (i.e., proportion of seen trials) for target (thick lines) and catch trials (thin lines) separately; Right, proportion of conscious trials (i.e., trials seen and correctly compared to 5); all as a function of target-mask delays for patients with OCD and good insight (blue), patients with OCD and poor insight (purple), patients with schizophrenia (pink) and healthy controls (black). The inset in the middle plane depicts detection *d*-prime, that is the ability of visibility ratings to distinguish between digit vs. catch trials. Thick lines correspond to fit of the logistic model. Error bars represent standard errors of the mean. **B**. Left, Schematic of the logistic model and its different free parameters (in gray). The consciousness threshold is determined by finding the target-mask delay for which 50% of the trials are consciously perceived (i.e., accurate and seen). Middle, Model parameters (i.e., upper asymptote, inflection point, lower asymptote, and slope) for each participant (dots) in each group. Horizontal lines represent the group averages. Overall, patients with OCD and good insight (blue) have similar parameter values as healthy controls (black), whereas patients with OCD and poor insight (purple) have similar parameter values as patients with schizophrenia (pink). Right, consciousness threshold measured on fitted logistic models in the different groups. **C**. Left, the logistic model was simulated with different upper asymptote values (between 50 and 100%, with 5% increments), all other parameters being held constant. Measured consciousness threshold (i.e., duration, in milliseconds, for which the logistic model equals 50%) decreases with increasing values of the upper asymptote. Middle, Proportion of conscious trials (as in panel A) before (dotted thin line) and after (thick line) correcting for non-one upper asymptotes on a participant basis. Right, consciousness threshold measured before (dotted lines) and after (thick lines) setting the upper asymptote to 1 in all groups. Group differences survive the correction for non-one upper asymptote.

### Higher threshold for conscious access in schizophrenia and OCD with poor insight

To more directly probe the integrity of conscious processing, we aggregated discrimination and visibility judgments and computed the proportion of conscious trials, which are trials in which the target digit was both accurately compared to 5 and rated as seen. The proportion of conscious trials confirmed results obtained with discrimination and visibility separately, with significant – and similar – impairments in patients with schizophrenia and patients with OCD and poor insight, but not in patients with OCD and good insight (**Supplementary Table 1**).

Measures of consciousness, such as the proportion of conscious trials, increase non-linearly with the target-mask delay, following a sigmoid function. This pattern of results is well accounted for by a logistic model whose parameters capture distinct aspects of non-conscious and conscious processing (**Fig. 2B**). We started by inspecting the consciousness threshold and found a significant difference across groups (controls: 52.8 ms, OCD with good insight: 47.5 ms, OCD with poor insight: 67.3 ms, schizophrenia: 73.2 ms; *F*_3,47_ = 7.61, *P <* 0.001, **Supplementary Table 2**). Notably, compared to healthy controls, consciousness threshold was elevated in patients with schizophrenia (*t*_21_ = 3.65, *P =* 0.002) and in patients with OCD and poor insight (*t*_10_ = 2.27, *P =* 0.048). By contrast, patients with OCD and good insight had a consciousness threshold significantly shorter to that of healthy controls (*t*_22_ = −2.15, *P =* 0.043). Moreover, there was no significant difference in consciousness thresholds between patients with OCD and poor insight, and patients with schizophrenia (*t*_18_ = −0.64, *P =* 0.53, 1/BF = 2.3). Crucially, in patients with OCD, there was a significant difference of consciousness threshold according to the level of insight (good versus poor insight: *t*_9_ = −3.15, *P =* 0.012, Cohen’s *d* = −1.5). In line with our main working hypothesis, these results demonstrate that the threshold for conscious access in OCD patients with poor insight more closely resemble that of patients with schizophrenia rather than that of OCD patients with good insight.

### Impairment of conscious processing in schizophrenia and OCD with poor insight

To study whether patients exhibit differences in conscious processing on top of elevated consciousness threshold, we compared the upper asymptotes of the fitted logistic model and found a significant difference across groups with the same pattern of significant impairments in patients with schizophrenia or with OCD and poor insight compared to healthy controls (controls: 95.4%, OCD with good insight: 96.3%, OCD with poor insight: 85.9%, schizophrenia: 84.1%; *F*_3,47_ = 5.33, *P =* 0.003, **Supplementary Table 2**). By contrast, there was no group difference for the lower asymptotes (*F*_3,47_ = 0.49, *P =* 0.69, 1/BF = 6.0). Altogether, this suggests that poor insight OCD and schizophrenia patients not only have a higher threshold for conscious access but also have impaired conscious processing.

### Impairment of conscious access and conscious processing co-exist in patients

Simulations showed that a decrease of the upper asymptotes mechanically decreases consciousness threshold and could therefore explain the pattern of results observed in patients with schizophrenia and OCD and poor insight (**Fig. 2C**). To address this concern, we corrected the data for non-one upper asymptotes on a participant-basis before re-fitting the logistic model. Expectedly, fitted upper asymptote parameters after correction were statistically indistinguishable from ideal ones (*t*-test against 1: all *P >* 0.2). Importantly, consciousness threshold similarly varied across groups after correction (controls: 52.1 ms, OCD with good insight: 47.0 ms, OCD with poor insight: 62.8 ms, schizophrenia: 66.5 ms; *F*_3,47_ = 6.64, *P <* 0.001) such that patients with schizophrenia still had an elevated consciousness threshold compared to healthy controls (*t*_23_ = −3.37, *P =* 0.003). Patients with OCD and poor insight had the same pattern of results but parametric statistics did not reach significance (*t*_10_ = 1.94, *P =* 0.081, BF = 2.52). Altogether, this analysis suggests that altered conscious processing and elevated consciousness threshold co-exist in schizophrenia and, to a lesser extent, in patients with OCD with poor insight.

### A shift in speed-accuracy tradeoff does not explain consciousness impairments

Impairment in conscious access and processing, as observed in patients with OCD and poor insight and in patients with schizophrenia, could result from a shift in speed-accuracy tradeoff whereby these patients would respond in haste and make more mistakes. To control for this possibility, we derived reaction time distributions separately for the discrimination and visibility tasks and estimated average reaction-times across groups (**Fig. 3A**). Patients with schizophrenia were significantly slower than any other group in performing the discrimination task (controls: 989 ms, OCD with good insight: 1010 ms, OCD with poor insight: 1134 ms, schizophrenia: 1414 ms, *F*_3,48_ = 10.62, *P <* 0.001) and in rating their visibility (controls: 694 ms, OCD with good insight: 673 ms, OCD with poor insight: 771 ms, schizophrenia: 886 ms, *F*_3,50_ = 4.70, *P =* 0.006, **Supplementary Table 1**). These findings thus preclude that differences between groups reflect shifted speed-accuracy tradeoff. On the one hand, patients with schizophrenia and patients with OCD and poor insight were slower, thus not explaining their lower discrimination and detection accuracy in terms of a speed-oriented tradeoff. On the other hand, OCD patients with good insight did not exhibit longer reaction times, that could have explained a lower consciousness threshold as an accuracy-oriented tradeoff.

**Fig. 3.**
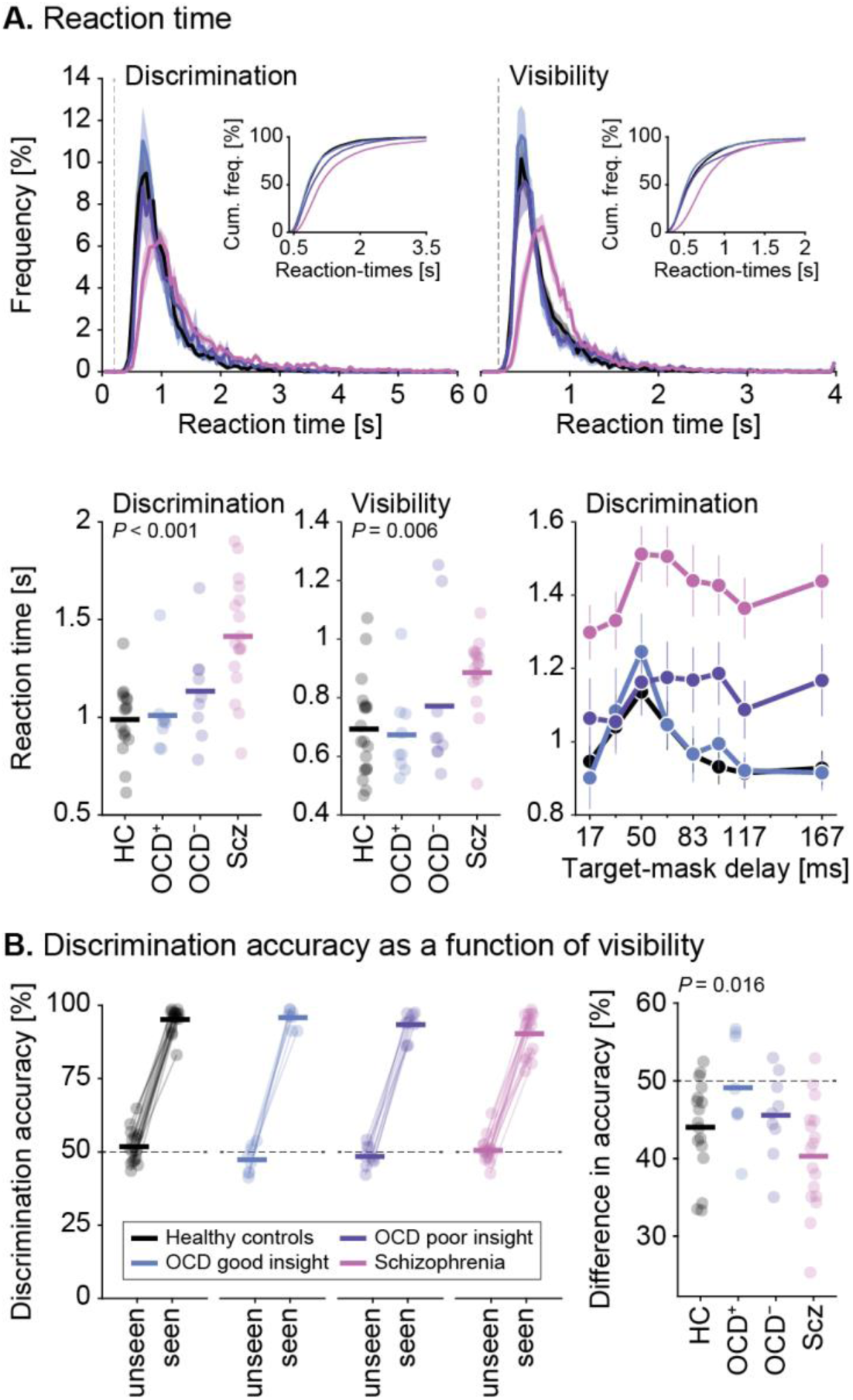
Non-consciousness related differences across groups. **A**. Top, Reaction time distributions corresponding to the discrimination and visibility questions. Insets depict the cumulative distributions. The vertical dotted gray line shows the lower cutoff for exclusion of abnormal reaction times. Bottom, Average reaction times corresponding to the discrimination and visibility questions; and discrimination reaction times as a function of target-mask delay. Patients with schizophrenia are significantly slower than any other group for both the discrimination and visibility tasks, thus precluding that differences in performance between groups are due to a shifted speed-accuracy tradeoff. **B**. Left, Discrimination accuracy is shown separately in target trials rated as seen vs. unseen. All participants have a discrimination accuracy at chance level for unseen trials and a near-ceiling accuracy for seen trials. Right, Sensitivity of visibility ratings to discrimination accuracy is measured by the difference in accuracy in unseen vs. seen trials. Patients with schizophrenia (pink) have seen vs. unseen difference in discrimination accuracy compared to the other groups (healthy controls, black, patients with OCD and good insight, blue, and with OCD and poor insight, purple). Each dot represents a participant and horizontal lines represents the group average.

We then inspected the evolution of reaction-times as a function of target-mask delay and observed that discrimination reaction times were significantly modulated by the target-mask delay (*F*_7,336_ = 10.03, *P <* 0.001). Moreover, there was an interaction between group and target-mask delay, whereby patients with OCD and poor insight appear to be slower compared to healthy controls (*F*_7,168_ = 3.85, *P <* 0.001) and patients with OCD and good insight (*F*_7,112_ = 3.94, *P <* 0.001) specifically at the long target-mask delays. We also observed that slower reaction-times occurred at intermediate delays where conscious perception is the most variable, splitting into conscious and non-conscious trials.^27^ Interestingly, the pattern of reaction-time variation was strikingly similar between good insight OCD patients and healthy controls on the one hand (with a narrow peak), and poor insight OCD and schizophrenia patients on the other hand (with a wide peak). By contrast, reaction times for the visibility question did not interact with the target-mask delay.

### Impairment of metacognitive sensitivity in schizophrenia

Since metacognitive ability in perceptual decision-making can be disturbed in patients with schizophrenia and OCD, we explored to what extent participants’ visibility ratings were predictive of discrimination accuracy in our task (**Fig. 3B**). As expected, discrimination accuracy for “unseen” trials, was close to chance level in all groups (controls: 52.5%, OCD with good insight: 48.0%, OCD with poor insight: 49.1%, schizophrenia: 51.2%; *t*-test against 50%: all *P >* 0.05), with no significant difference between groups (*F*_3,47_ = 1.96, *P =* 0.13, 1/BF = 1.5).

To quantify the correspondence between objective decisions and subjective reports, we computed the difference in discrimination accuracy between seen and unseen trials. This measure significantly differed across groups (controls: 44.0%, OCD with good insight: 49.2%, OCD with poor insight: 45.6%, schizophrenia: 40.3%, *F*_3,47_ = 3.78, *P =* 0.016, **Supplementary Table 1**). In particular, seen vs. unseen difference in discrimination accuracy was lower for patients with schizophrenia compared to patients with OCD (good insight: *t*_15_ = −3.05, *P =* 0.008; poor insight: *t*_20_ = −2.10, *P =* 0.049).

### PCA decomposition of clinical scales reveals meaningful clinical dimensions

To reveal clinical dimensions beyond classical nosography as well as to reduce symptom dimensionality, we applied principal component analysis (PCA) on all items from relevant clinical scales in patient groups (Y-BOCS, OVIS, HAM-A, HAM-D and DOCS scales in patients with OCD and SANS, SAPS and CDS in patients with schizophrenia).

In the group of patients with OCD, PCA decomposition of clinical scales yielded 3 significant principal components (PC) accounting for 53.7% of the symptoms (**Fig. 4A**). The first PC (PC1), represents global psychopathology, with almost all items having positive loadings. By contrast, PC2 is dominated by OVIS items. Interestingly, other loadings suggest that patients with poor insight have intense cleaning symptoms but less unacceptable thoughts obsessions and somatic symptoms compared to patients with good insight. Finally, PC3 reveals an inverse relationship between contamination and responsibility concerns. Importantly, this PC also suggests that contamination symptoms are more associated with suicide than unacceptable thoughts obsessions.

**Fig. 4.**
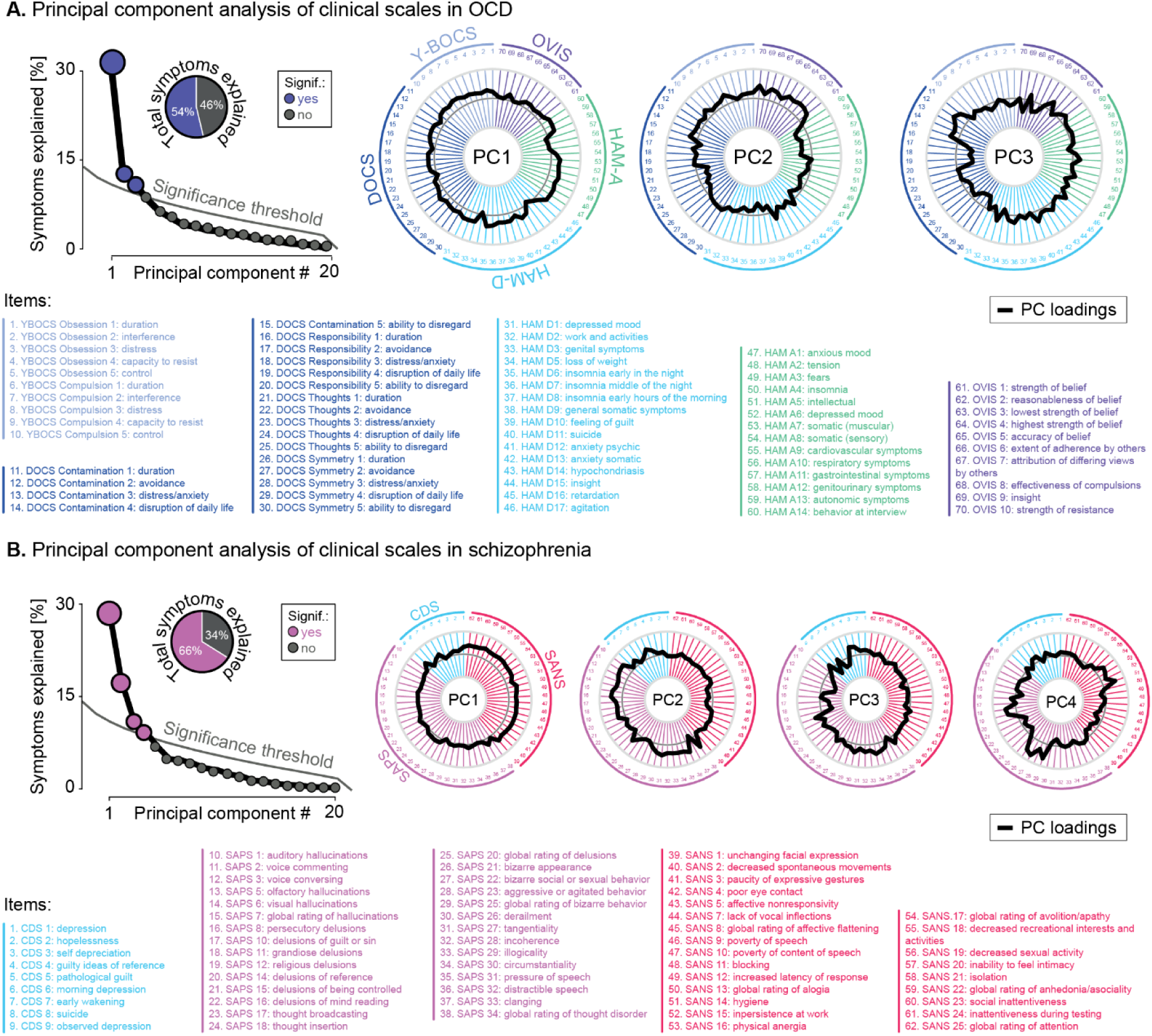
Clinical dimensions yielded by principal component analysis. **A**. In the group of patients with OCD, PCA yielded 3 significant principal components accounting for the 55% of the symptoms. In PC1, accounting for 31.4% of the variance, almost all items have positive loadings except one item of the DOCS (avoidance related to unpleasant thoughts), some physical items of the HAM-D (hypochondriasis, loss of weight) and of the HAM-A (cardiovascular and respiratory symptoms). PC2, accounting for 12.9% of the variance, was dominated by the OVIS items and also shows high loadings for Y-BOCS, the contamination items of the DOCS and suicide, but negative loadings for the thought items of the DOCS and many items of HAM-A and HAM-D (HAM-A: somatic symptoms, cardiovascular, genitourinary, autonomic symptoms, HAM-D: retardation, insomnia, feeling of guilt). PC3, accounting for 10.9% of the variance, shows an inverse relationship between contamination concerns, on the one hand, and the responsibility concerns (and to a lesser extent, symmetry concerns) on the other hand, as measured by the DOCS. **B**. In the group of patients with schizophrenia, PCA yielded 4 significant PC accounting for 66% of the symptoms. In PC1, accounting for 32.4% of the variance, all SANS items have high loadings, as well as several items of the CDS (“depression,” “morning depression,” “observed depression”). PC2, accounting for 20.2% of the variance, is highly dominated by positive symptoms (all items of hallucinations, all items of thought disorder, except circumstantiality, many delusional items apart from influence syndrome) and few depressive symptoms (pathological guilt and suicide). Regarding negative symptoms, there was a negative loading for several alogia symptoms. PC3, accounting for 11.6% of the variance, has positive and negative loadings both in the positive symptoms and the depression dimension. More specifically, it shows an anticorrelation between, on the one hand, delusion of reference, most of the items related to thought disorder in the SAPS and guilty ideas of reference and self-depreciation in the CDS and, on the other hand, hallucinations (visual, olfactory and voices), delusions (religious, guilt, grandiose) and specific symptoms of depression (suicide, pathological guilt and morning depression). PC4 accounting for 10.0% of the variance, captures mostly bizarre behavior, and a mixture of positive and negative symptoms (religious and grandiose delusions, avolition including hygiene and apathy).

In the group of patients with schizophrenia, PCA yielded 4 significant PC accounting for 66% of the symptoms (**Fig. 4B**). PC1 mostly reflects negative symptoms, as well as specific depressive symptoms. PC2 is highly dominated by positive symptoms and few depressive symptoms. PC3 captures a most complex pattern, characterized by positive and negative loadings, both in the positive symptoms and the depression dimensions. More specifically, it shows an anticorrelation between, on the one hand, delusion of reference, most of the items related to thought disorder and specific items of the CDS and on the other hand, hallucinations, delusions and other symptoms of depression. PC4 captures mostly bizarre behavior, and a mixture of delusional and avolition symptoms.

### Age correlates with consciousness threshold

We then investigated to what extent disruption of consciousness related to clinical variables. To do so, we first checked whether general socio-demographic factors (i.e., age, gender, smoking) would influence consciousness measures across participants. We found a significant effect of age on most of the consciousness measures (consciousness threshold, discrimination accuracy, detection *d*-primes, discrimination and visibility thresholds, reaction times for both tasks, all *P <* 0.05), older people having on average more conscious impairments. There was no interaction with the group and no effect of gender or of the smoking status (all *P >* 0.2). No interaction was found between age, gender or smoking status and upper asymptotes, or seen vs. unseen difference in discrimination accuracy (all *P >* 0.05).

### Insight correlates with consciousness threshold in OCD patients

We then run an ANOVA to measure how consciousness measures were explained by clinical variables with the target-mask delay and age as covariates whenever relevant. We first confirmed the significant interaction between OVIS and consciousness threshold in the OCD group (*F*_1,14_ = 9.22, *P =* 0.009, **Fig. 5A**). We did not find any other significant interaction (all *P >* 0.1), notably no significant interaction with fluoxetine equivalent (*P =* 0.86, 1/BF = 2.4). Regarding PC, only PC2 significantly interacted with the consciousness threshold (*F*_1,14_ = 9.78, *P =* 0.007; for other PC: *P >* 0.8), the upper asymptote, and all other measures of consciousness such as discrimination and visibility thresholds, discrimination accuracy, detection *d*-primes (all *P <* 0.05).

**Fig. 5.**
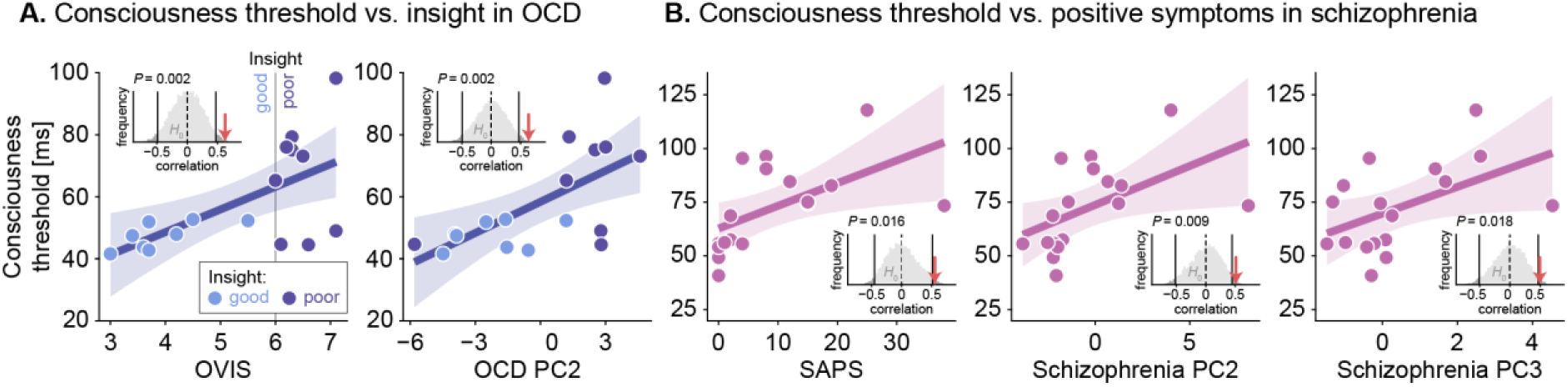
Correlation between consciousness threshold and clinical measures. **A**. In the group of patients with OCD, consciousness thresholds (in ms) significantly correlate with OVIS and PC2 scores (which included insight items). The vertical gray line represents the OVIS threshold used to distinguish patients with good (OVIS < 6, blue color) and poor insight (OVIS ≥ 6, purple color). **B**. In the group of patients with schizophrenia (pink), consciousness thresholds significantly correlate with SAPS, PC2 scores (which included hallucinations and delusions items) and PC3 scores (which includes both positive and depressive symptoms, including delusion of reference, thought disorder, guilty ideas of reference and self-depreciation). Each dot represents a participant, lines represent linear regressions, and shaded areas represent 95% confidence intervals. The bar plots represent null distributions of the correlation coefficient between each clinical measure and consciousness threshold (10,000 permutations), significance thresholds (vertical black lines), and the observed correlation value (red arrow).

The seen vs. unseen difference in discrimination accuracy did not significantly interact with OVIS or PC2 (all *P >* 0.1), but was negatively associated with the Y-BOCS total scores (*F*_1,14_ = 5.69, *P =* 0.032, **Supplementary Fig. 2A**) and its obsession subscore (*F*_1,14_ = 9.22, *P =* 0.009): patients with higher levels of obsessions had less difference between seen and unseen trials. When splitting OCD group according to insight, we observed that this pattern was driven by patients with poor insight (Y-BOCS total scores: *F*_1,6_ = 7.22, *P =* 0.036; Y-BOCS obsession score: *F*_1,6_ = 10.07, *P =* 0.019), whereas patients with OCD and good insight had a strong effect of the HAM-A (*F*_1,6_ = 17.01, *P =* 0.009) and an effect of fluoxetine doses (*F*_1,6_ = 6.70, *P =* 0.049) instead.

Reaction times for the discrimination task also significantly interacted with PC2 (*F*_1,15_ = 5.66, *P =* 0.031), but not with OVIS (*F*_1,15_ = 2.02, *P =* 0.18, 1/BF = 1.2). Moreover, they positively interacted with the Y-BOCS (total scores: *F*_1,15_ = 6.14, *P =* 0.026, obsession scores: *F*_1,15_ = 5.96, *P =* 0.027, compulsion scores: *F*_1,15_ = 5.22, *P =* 0.037, **Supplementary Fig. 2A**) and the DOCS symmetry scores (*F*_1,15_ = 7.85, *P =* 0.013). When splitting OCD group according to insight, we found that the association between obsessions and discrimination reaction times was specific to patients with good insight (Y-BOCS total scores: *F*_1,6_ = 7.37, *P =* 0.035, Y-BOCS compulsion scores: *F*_1,6_ = 6.52, *P =* 0.043; DOCS total: *F*_1,6_ = 7.92, *P =* 0.031; DOCS symmetry scores: *F*_1,6_ = 12.83, *P =* 0.012; Y-BOCS obsession scores: *F*_1,6_ = 5.54, *P =* 0.057) with PC1 being significant (*F*_1,6_ = 11.24, *P =* 0.015), all results being non-significant in the poor insight group (all *P >* 0.05).

No effect was observed for reaction times in the visibility task.

### Positive symptoms correlate with consciousness threshold in schizophrenia

Running the same ANOVAs in the group of patients with schizophrenia, we found that SAPS scores significantly interacted with consciousness threshold (*F*_1,14_ = 6.38, *P =* 0.024, **Fig. 5B**) without any interaction with the SANS, with the CDS scores (all *P >* 0.1), and with olanzapine equivalent (*P =* 0.71, 1/BF = 2.3). Regarding PC, we found that PC2 and PC3 significantly interacted with the consciousness threshold (PC2: *F*_1,14_ = 5.35, *P =* 0.037; PC3: *F*_1,14_ = 4.63, *P =* 0.049; others PC: *P >* 0.4).

Discrimination accuracy significantly interacted with SAPS (*F*_1,14_ = 8.00, *P =* 0.013) and PC2 (*F*_1,14_ = 6.18, *P =* 0.026), whereas discrimination threshold only correlated with SAPS (*F*_1,14_ = 5.12, *P =* 0.040). Upper asymptotes, detection *d*-primes, visibility threshold and seen vs. unseen difference in discrimination accuracy did not significantly interact with clinical measures (all *P >* 0.05).

For the discrimination reaction times, we observed strong interactions between target-mask delays and several clinical measures (SAPS, SANS, PC1, PC2, all *P <* 0.01), suggesting that these measures were influenced by different clinical variables at short and long target-mask delays. To further understand these results, we conducted post-hoc exploratory analyses, by looking at those interactions separately for short and long target-mask delays (3 shortest, i.e., 16.7, 33.4, and 50.1 ms, on the one hand, and 3 longest, i.e., 100.2, 116.9, 167.0 ms, on the other hand). We found that reaction times were modulated by SANS and PC1 at the shortest target-mask delays (*F*_1,14_ = 5.14, *P =* 0.040 and *F*_1,14_ = 5.11, *P =* 0.040 respectively, **Supplementary Fig. 2B**) whereas they were modulated by the SAPS and the PC2 at the longest target-mask delays (*F*_1,14_ = 8.63, *P =* 0.011 and *F*_1,14_ = 8.34, *P =* 0.012 respectively). No effect was observed for visibility reaction times.

## DISCUSSION

### Summary of the results

In this study, we found that conscious access and conscious processing were both impaired in patients with schizophrenia and in patients with OCD who had a poor insight. By contrast, patients with OCD and good insight had a lower consciousness threshold than healthy controls. A PCA decomposition of clinical scales revealed meaningful clinical dimensions within each group, capturing how symptoms covary across distinct clinical scales. In patients with schizophrenia, impairments of consciousness were associated with positive symptoms as assessed by the SAPS and relevant PCs. In the OCD group, impairments of consciousness were related to the OVIS scores and the corresponding clinical dimension computed with PCA.

### Disruption of conscious access in schizophrenia is linked to positive symptoms

In the schizophrenia group, our results are in line with previous findings, showing that those patients have an impaired conscious access and abnormal conscious processing, whereas subliminal processing is preserved.^2^ Interestingly, disruption of conscious access could favor the advent of psychotic symptoms.^4^ However, few previous studies examined the relationship between clinical dimensions and consciousness disruption, and with global clinical scores only.^3,4,22,33^ The only study which included clinical subscales found no differential impact of subscores on consciousness impairments,^39^ possibly because patients were less symptomatic or explored with less sensitive clinical scales. In the current study, disruption of consciousness was associated with different aspects of positive symptoms. All positive symptoms including PC2 (that is dominated by delusional items) and PC3 (which captures positive and depressive dimensions) correlated with consciousness thresholds. By contrast, discrimination measures mostly depended on delusional symptoms and thought disorder independently from depressive symptoms (SAPS and PC2 but not PC3) and conscious processing (upper asymptotes) and visibility were not significantly impacted by any clinical measure.

We also found that patients with schizophrenia had longer reaction times when performing the discrimination task. More specifically, negative symptoms tended to slow down responses for short target-mask delays (subliminal processing), whereas positive symptoms were associated with delayed responses for longer target-mask delays. This could reflect that participants with many positive symptoms needed more time to process conscious information, therefore constituting an indirect sign of conscious processing impairment. In line with this hypothesis, patients with OCD and poor insight, who also exhibit consciousness disruption in our study, appeared to be specifically slowed down for long target-mask delays.

We previously proposed that conscious access disruption enhanced adherence to beliefs disconnected from the reality, which may culminate in delusions.^2^ Indeed, when access to external information is reduced, patients tend to be interpretative, and beliefs constructed from this sparse information could be subsequently strengthened by a bias favoring conscious access to confirmatory evidence. On top of those alterations, abnormal attentional amplification^22^ could orient conscious access to random rather than relevant external information, giving rise to a feeling of strangeness regarding consciousness content and, in extreme cases, to an impression of xenopathy.

In the current study, we also found that impaired conscious access was associated with the advent of hallucinations, which have high loadings in PC2. In line with previous proposals,^2,40,41^ this result emphasizes that hallucinations are internal constructions likely favored by a reduced sensitivity to the external world.

### Consciousness threshold, obsessions and insight

Importantly, our results confirm that an impairment of consciousness does not necessarily lead to psychotic symptoms.^5–7^ Indeed, we demonstrate for the first time that consciousness is disrupted in a specific subgroup of patients with OCD, characterized by poor insight, with a profile similar to that of patients with schizophrenia for most measures. By contrast, patients with OCD and good insight have similar or even better consciousness measures compared to healthy controls.

Previous studies showed that patients with OCD exhibit both a reduced cognitive inhibition and a deficit in selective attention making them less able to reject distractors and intrusive thoughts.^42–45^ In this sense, they could have a decreased, rather than an increased, consciousness threshold for irrelevant information, especially for internal representations and intrusive thoughts. Our task was not specifically designed to examine internal thoughts. However, our results suggest that in OCD, conscious access to the external world is normal – or perhaps even slightly enhanced – when insight is preserved, allowing them to be critical about their obsessions, whereas it is on the contrary decreased in case of poor insight. One interesting possibility is that a limited access to external stimulation, as measured in our experiment, could leave more room for internal representations that could degenerate into obsessions. Under these circumstances, patients may have more difficulty to perceive the pathological nature of their symptoms as their thought content would only be shifted from external stimulation towards internal representations and has therefore no reason to be egodystonic.

We also observed that obsession levels had a negative impact on discrimination reaction times and decreased the seen vs. unseen difference in discrimination accuracy. Interestingly, those effects differed in patients with good vs. poor insight. In OCD patients with good insight, obsession scales modulated reaction times but not difference in accuracy (which were rather dependent of anxiety levels). By contrast, obsession scales modulated difference in accuracy but not reaction times in OCD patients with poor insight. As it has been shown that obsessions interfere with decision-making and alter confidence,^46–48^ a plausible explanation is that patients with good insight and high level of obsession may take more time in order to feel more confident,^49–51^ and may more easily rate a trial as “unseen” whenever unsure, with a preserved (or improved) accuracy. By contrast, patients with poor insight and high level of obsession may not specifically compensate their level of uncertainty, so that the interferences due to obsessions result in a decrease in discrimination accuracy for “seen” trials, and ultimately a smaller seen vs. unseen difference in discrimination accuracy.^8^ Overall, those differences highlight that patients with OCD with good and poor insight may have distinct neuropsychological functioning and adaptation to obsessive-compulsive symptoms.

### From lack of insight to delusions

One naturally arising question is, if consciousness is disrupted in both patients with schizophrenia and patients with poor insight OCD, why this would participate to the advent of psychotic symptoms in schizophrenia but not in OCD. In search of an answer, we have looked at consciousness impairments that would be specifically found in schizophrenia. One of the only differences that we observed between patients with OCD and poor insight and patients with schizophrenia (on top of slower reaction times for the discrimination task) is of metacognitive nature, whereby the difference in discrimination accuracy between seen and unseen trials was weaker in schizophrenia. In other words, patients with OCD and poor insight are less able to consciously perceive external information whereas patients with schizophrenia have a double deficit: they perceive less information and tend to make more mistakes when they consider to have properly seen the information. This second deficit can be interpreted in three different and non-exclusive ways. First, patients with schizophrenia may be overconfident in their ability to see and rate as “seen” information that they in fact did not consciously perceive. Alternatively, they may be less impaired in detection than in discrimination, resulting in situations where they have indeed seen something but are not able to identify it. Finally, they may sometimes “hallucinate” rather than see the target stimulus, resulting in a misclassification (but note, however, that they do not exhibit more false alarms for catch trials). Even if the seen vs. unseen difference in discrimination accuracy and visibility were not significantly associated with positive symptoms, we found that those symptoms were highly correlated with a decrease in discrimination, a result that is compatible with a dissociation between detection and discrimination. Irrespective of its etiology, such a discrepancy between mental experience and external events may destabilize trust in others. Indeed, a rational explanation of being contradicted by others when overly confident is that others’ reports are wrong or deliberate lies. Similarly, having the impression to see information that is difficult to discriminate likely increases proneness to interpretations. All in all, these abnormalities may fuel delusional constructions with persecution content (which have high loadings in PC2).

### Patients with OCD and poor insight: closer to schizophrenia than to OCD?

Patients with OCD and poor insight have been shown to exhibit more severe illness, more obsessive-compulsive symptoms, consistent with our results (PC2 shows positive loadings for symptom severity in the Y-BOCS), as well as lower response to treatment.^52–54^ Moreover, they share neuropsychological deficits and neural overlaps with patients with schizophrenia.^8,11–13,55–57^ They also have higher rates of schizophrenia spectrum disorders in their first-degree relatives.^58^ In our study, they strikingly resemble patients with schizophrenia in terms of consciousness disruption while they strongly differ from OCD patients with good insight. Their consciousness deficit could be underpinned by neuronal features common with schizophrenia, such as a decrease of the event-related potential P300^59,60^ or large-scale brain dysconnectivity.^61–63^ Overall, our results support the hypothesis of a continuum between OCD and schizophrenia, including patients having an OCD with poor insight, or with psychotic features, and a schizo-obsessive subtype of schizophrenia (i.e., patients with schizophrenia having obsessive-compulsive symptoms).^58,64–67^ Interestingly, some patients with refractory OCD benefit from an augmentation of their treatment by an antipsychotic.^68,69^ In our cohort, no OCD patient was treated with antipsychotics precluding any analysis of their potential impact on consciousness disruption. Further explorations could determine whether patients who have impairments of consciousness and poor insight are more likely to respond to antipsychotics, allowing a better adjustment of therapeutic strategies to patients’ cognitive profile.

### Limitations

The main limitation of our study is that we did not include other patient groups exhibiting a lack of insight, or use a specific scale to measure insight in the schizophrenia group, so we could not explore whether the relationship between consciousness impairment and insight was restricted to patients with OCD. Our sample size is also relatively small, some analyses may thus lack power.

### Conclusions

Overall, our study provides new insights about consciousness disruption in psychiatric disorders. It confirms that impairment of consciousness is observed in patients with schizophrenia and associated with positive symptoms, but also demonstrates that it can manifest as a lack of insight in patients with OCD, delineating a distinct subgroup in terms of cognitive deficits and clinical features. Subdividing psychiatric symptomatology into dimensions associated with neurocognitive profiles is a challenge, notably in OCD.^70–80^ Using PCA, we were able to obtain data-driven clinical descriptions, illustrating how symptoms interact beyond classical clinical scales. We show that specific symptoms correlate to consciousness disruption, thereby refining the mapping between semiology and cognition. Further studies are needed to confirm these results, extend these findings to other clinical populations with good and poor insight, better characterize the neurocognitive mechanisms involved in the disruption of consciousness in psychiatric disorders and assess whether it should benefit from specific therapeutics.

## DATA AVAILABILITY

Custom codes written in *R* and *MATLAB* are available from the corresponding author upon reasonable request.

## ACKNOWLEDGEMENTS

We thank all the participants. LB thanks the *Fondation Bettencourt-Schueller*, the *Philippe Foundation,* the *Foundation L’Oréal-Unesco* and the National Institute of Mental Health (R01MH116038 and U01MH121766) for their support. MM thanks the *Alexander von Humboldt Stiftung* and the *Fondation Bettencourt-Schueller* for their support.

## AUTHOR CONTRIBUTIONS

ST, BY and MG had the initial idea; LB coded the experiment; ST, BY and MG collected the data; LB and MM designed the methodology for the analyses; LB performed the analyses; LB wrote the initial draft of the manuscript; MM made the figures; MM, ST, BY and MG reviewed and edited the manuscript.

