## Supplemental materials for "Disruption of consciousness depends on insight in OCD and on positive symptoms in schizophrenia"

### Supplementary table 1. Consciousness measures

| Measures | *F*-value | *p*-value | Bayes Factor |
|---|---|---|---|
| **Discrimination** | | | |
| **Target-mask delay** | $\mathbf{F_{7,336} = 361.91}$ | $\mathbf{P < 0.001}$ | – |
| **Group** | $\mathbf{F_{3,48} = 8.66}$ | $\mathbf{P < 0.001}$ | – |
| **Group × target-mask delay** | $\mathbf{F_{21,336} = 5.11}$ | $\mathbf{P < 0.001}$ | – |
| **Schizophrenia vs. healthy controls** | | | |
| Group | $\mathbf{F_{1,32} = 14.75}$ | $\mathbf{P < 0.001}$ | – |
| Group × target-mask delay | $\mathbf{F_{7,224} = 5.68}$ | $\mathbf{P < 0.001}$ | – |
| **OCD poor insight vs. healthy controls** | | | |
| Group | $\mathbf{F_{1,24} = 11.51}$ | $\mathbf{P = 0.002}$ | – |
| Group × target-mask delay | $\mathbf{F_{7,168} = 6.22}$ | $\mathbf{P < 0.001}$ | – |
| **OCD good insight vs. healthy controls** | | | |
| Group | $F_{1,24} = 0.27$ | $P = 0.61$ | 1/BF = 6.1 |
| Group × target-mask delay | $F_{7,168} = 1.55$ | $P = 0.16$ | 1/BF = 5.8 |
| **OCD poor insight vs. schizophrenia** | | | |
| Group | $F_{1,24} = 0.10$ | $P = 0.76$ | 1/BF = 5.9 |
| Group × target-mask delay | $F_{7,168} = 0.66$ | $P = 0.71$ | 1/BF = 26.5 |
| **OCD good vs. poor insight** | | | |
| Group | $\mathbf{F_{1,16} = 12.50}$ | $\mathbf{P = 0.003}$ | – |
| Group × target-mask delay | $\mathbf{F_{7,112} = 10.28}$ | $\mathbf{P < 0.001}$ | – |
| **Visibility for non-catch trials[1]** | | | |
| **Target-mask delay** | $\mathbf{F_{7,350} = 580.07}$ | $\mathbf{P < 0.001}$ | – |
| **Group** | $\mathbf{F_{3,50} = 5.56}$ | $\mathbf{P = 0.002}$ | – |
| **Group × target-mask delay** | $\mathbf{F_{21,350} = 3.91}$ | $\mathbf{P < 0.001}$ | – |
| **Schizophrenia vs. healthy controls** | | | |
| Group | $\mathbf{F_{1,34} = 11.88}$ | $\mathbf{P = 0.002}$ | – |
| Group × target-mask delay | $\mathbf{F_{7,238} = 5.2}$ | $\mathbf{P < 0.001}$ | – |
| **OCD poor insight vs. healthy controls** | | | |
| Group | $\mathbf{F_{1,26} = 7.75}$ | $\mathbf{P = 0.010}$ | **–** |
| Group × target-mask delay | $\mathbf{F_{7,182} = 3.59}$ | $\mathbf{P = 0.001}$ | **–** |
| **OCD good insight vs. healthy controls** | | | |
| Group | $F_{1,26} = 0.16$ | $P = 0.69$ | 1/BF = 6.5 |
| Group × target-mask delay | $\mathbf{F_{7,182} = 2.06}$ | $\mathbf{P = 0.049}$ | – |
| **OCD poor insight vs. schizophrenia** | | | |
| Group | $F_{1,24} = 0.076$ | $P = 0.79$ | 1/BF = 6.1 |

| Measures | *F*-value | *p*-value | Bayes Factor |
|---|---|---|---|
| Group × target-mask delay | $F_{7,168}$ = 0.22 | *P* = 0.98 | 1/BF = 31.9 |
| **OCD good vs. poor insight** | | | |
| Group | **$F_{1,16}$ = 4.80** | ***P* = 0.044** | – |
| Group × target-mask delay | **$F_{7,112}$ = 5.65** | ***P* < 0.001** | – |
| **Detection *d*-primes** | | | |
| **Target-mask delay** | **$F_{7,350}$ = 190.15** | ***P* < 0.001** | **–** |
| **Group** | **$F_{3,50}$ = 5.66** | ***P* = 0.002** | **–** |
| **Group × target-mask delay** | $F_{21,350}$ = 1.04 | *P* = 0.41 | 1/BF = 116.2 |
| **Schizophrenia vs. healthy controls** | | | |
| Group | **$F_{1,34}$ = 7.11** | ***P* = 0.012** | – |
| Group × target-mask delay | $F_{7,238}$ = 1.16 | *P* = 0.33 | 1/BF = 18.9 |
| **OCD poor insight vs. healthy controls** | | | |
| Group | **$F_{1,26}$ = 11.04** | ***P* = 0.003** | – |
| Group × target-mask delay | $F_{7,182}$ = 0.28 | *P* = 0.96 | 1/BF = 31.6 |
| **OCD good insight vs. healthy controls** | | | |
| Group | $F_{1,26}$ = 0.41 | *P* = 0.53 | 1/BF = 5.4 |
| Group × target-mask delay | $F_{7,182}$ = 0.56 | *P* = 0.79 | 1/BF = 23.5 |
| **OCD poor insight vs. schizophrenia** | | | |
| Group | $F_{1,24}$ = 0.42 | *P* = 0.52 | 1/BF = 4.9 |
| Group × target-mask delay | $F_{7,168}$ = 0.95 | *P* = 0.47 | 1/BF = 14.5 |
| **OCD good vs. poor insight** | | | |
| Group | **$F_{1,16}$ = 10.86** | ***P* = 0.005** | – |
| Group × target-mask delay | $F_{7,112}$ = 0.83 | *P* = 0.56 | 1/BF = 12.6 |
| **Proportion of conscious trials** | | | |
| **Target-mask delay** | **$F_{7,329}$ = 607.64** | ***P* < 0.001** | – |
| **Group** | **$F_{3,47}$ = 9.64** | ***P* < 0.001** | – |
| **Group × target-mask delay** | **$F_{21,329}$ = 5.43** | ***P* < 0.001** | – |
| **Schizophrenia vs. healthy controls** | | | |
| Group | **$F_{1,32}$ = 16.48** | ***P* < 0.001** | – |
| Group × target-mask delay | **$F_{7,224}$ = 6.77** | ***P* < 0.001** | – |
| **OCD poor insight vs. healthy controls** | | | |
| Group | **$F_{1,24}$ = 11.32** | ***P* = 0.003** | – |
| Group × target-mask delay | **$F_{7,168}$ = 5.27** | ***P* < 0.001** | – |
| **OCD good insight vs. healthy controls** | | | |
| Group | $F_{1,23}$ = 2.51 | *P* = 0.13 | 1/BF = 4.8 |
| Group × target-mask delay | **$F_{7,161}$ = 2.61** | ***P* = 0.014** | – |
| **OCD poor insight vs. schizophrenia** | | | |

| Measures | *F*-value | *p*-value | Bayes Factor |
|---|---|---|---|
| Group | $F_{1,24} = 0.22$ | $P = 0.64$ | 1/BF = 5.6 |
| Group × target-mask delay | $F_{7,168} = 0.24$ | $P = 0.98$ | 1/BF = 35.0 |
| **OCD good vs. poor insight** | | | |
| Group | $\mathbf{F_{1,15} = 12.27}$ | $\mathbf{P = 0.003}$ | – |
| Group × target-mask delay | $\mathbf{F_{7,105} = 10.06}$ | $\mathbf{P < 0.001}$ | – |
| **Difference of discrimination accuracy between seen and unseen trials** | | | |
| **Group** | $\mathbf{F_{3,47} = 3.78}$ | $\mathbf{P = 0.017}$ | – |
| Schizophrenia vs. healthy controls | $t_{31} = -1.65$ | $P = 0.11$ | BF = 1.1 |
| **OCD poor insight vs. healthy controls** | $t_{18} = 0.66$ | $P = 0.52$ | 1/BF = 2.7 |
| **OCD good insight vs. healthy controls** | $t_{13} = 1.85$ | $P = 0.088$ | BF = 1.4 |
| **Schizophrenia vs. OCD good insight** | $\mathbf{t_{15} = -3.05}$ | $\mathbf{P = 0.008}$ | – |
| **OCD poor insight vs. schizophrenia** | $\mathbf{t_{20} = 2.10}$ | $\mathbf{P = 0.049}$ | – |
| **OCD good vs. poor insight** | $t_{14} = 1.20$ | $P = 0.25$ | 1/BF = 1.5 |
| **OCD poor insight vs. healthy controls** | $t_{18} = 0.66$ | $P = 0.52$ | 1/BF = 2.7 |
| **Discrimination reaction times** | | | |
| **Target-mask delay** | $\mathbf{F_{7,336} = 10.03}$ | $\mathbf{P < 0.001}$ | – |
| **Group** | $\mathbf{F_{3,48} = 10.65}$ | $\mathbf{P < 0.001}$ | – |
| **Group × target-mask delay** | $\mathbf{F_{21,336} = 1.73}$ | $\mathbf{P = 0.026}$ | – |
| **Schizophrenia vs. healthy controls** | | | |
| Group | $\mathbf{F_{1,32} = 26.17}$ | $\mathbf{P < 0.001}$ | – |
| Group × target-mask delay | $F_{7,224} = 1.93$ | $P = 0.066$ | 1/BF = 23.1 |
| **OCD poor insight vs. healthy controls** | | | |
| Group | $F_{1,24} = 2.93$ | $P = 0.10$ | BF = 1.1 |
| Group × target-mask delay | $\mathbf{F_{7,168} = 3.85}$ | $\mathbf{P < 0.001}$ | – |
| **OCD good insight vs. healthy controls** | | | |
| Group | $F_{1,24} = 0.08$ | $P = 0.78$ | 1/BF = 5.2 |
| Group × target-mask delay | $F_{7,168} = 0.98$ | $P = 0.45$ | 1/BF = 31.1 |
| **OCD poor insight vs. schizophrenia** | | | |
| Group | $\mathbf{F_{1,24} = 5.92}$ | $\mathbf{P = 0.023}$ | **–** |
| Group × target-mask delay | $F_{7,168} = 0.29$ | $P = 0.96$ | 1/BF = 34.3 |
| **Schizophrenia vs. OCD good insight** | | | |
| Group | $\mathbf{F_{1,24} = 13.56}$ | $\mathbf{P = 0.001}$ | **–** |
| Group × target-mask delay | $F_{7,168} = 1.67$ | $P = 0.11$ | 1/BF = 35.9 |
| **OCD good vs. poor insight** | | | |
| Group | $F_{1,16} = 1.32$ | $P = 0.27$ | 1/BF = 1.6 |
| Group × target-mask delay | $\mathbf{F_{7,112} = 3.94}$ | $\mathbf{P < 0.001}$ | – |

| Measures | $F$-value | $p$-value | Bayes Factor |
|---|---|---|---|
| **Visibility reaction times** | | | |
| **Target-mask delay** | $F_{7,350} = 1.62$ | $P = 0.13$ | 1/BF = 496.4 |
| **Group** | **$F_{3,50} = 4.70$** | **$P = 0.006$** | – |
| **Group × target-mask delay** | $F_{21,350} = 1.02$ | $P = 0.43$ | 1/BF = 843.2 |
| **Schizophrenia vs. healthy controls** | **$t_{33} = -3.94$** | **$P < 0.001$** | – |
| **OCD poor insight vs. healthy controls** | $t_{11} = -0.81$ | $P = 0.43$ | BF = 1.9 |
| **OCD good insight vs. healthy controls** | $t_{17} = 0.32$ | $P = 0.76$ | 1/BF = 2.6 |
| **Schizophrenia vs. OCD good insight** | **$t_{14} = -3.59$** | **$P = 0.003$** | – |
| **OCD poor insight vs. schizophrenia** | $t_{10} = 1.23$ | $P = 0.25$ | 1/BF = 1.2 |
| **OCD good vs. poor insight** | $t_{13} = 0.97$ | $P = 0.35$ | 1/BF = 1.8 |

[1] No significant difference was observed in the number of false alarms among the catch trials (all $P > 0.1$ and 1/BF > 5).

### Supplementary table 2. Parameters of logistic model

| Measures | *F*-value | *p*-value | Bayes Factor |
|---|---|---|---|
| **Discrimination threshold** | | | |
| **Group** | $F_{3,48} = 6.98$ | $P < 0.001$ | – |
| **Schizophrenia vs. healthy controls** | $t_{23} = 3.42$ | $P = 0.002$ | – |
| **OCD poor insight vs. healthy controls** | $t_{11} = 2.98$ | $P = 0.012$ | – |
| **OCD good vs. poor insight** | $t_{10} = -3.32$ | $P = 0.008$ | – |
| **OCD good insight vs. healthy controls** | $t_{24} = -0.45$ | $P = 0.66$ | 1/BF = 2.5 |
| **OCD poor insight vs. schizophrenia** | $t_{20} = -0.21$ | $P = 0.84$ | 1/BF = 2.6 |
| **Visibility threshold** | | | |
| **Group** | $F_{3,50} = 5.13$ | $P = 0.004$ | – |
| **Schizophrenia vs. healthy controls** | $t_{24} = 3.17$ | $P = 0.005$ | – |
| **OCD poor insight vs. healthy controls** | $t_{10} = 1.82$ | $P = 0.097$ | BF = 2.1 |
| **OCD good vs. poor insight** | $t_{14} = -2.03$ | $P = 0.062$ | BF = 1.6 |
| **OCD good insight vs. healthy controls** | $t_{14} = -0.63$ | $P = 0.54$ | 1/BF = 2.3 |
| **OCD poor insight vs. schizophrenia** | $t_{17} = 0.59$ | $P = 0.56$ | 1/BF = 2.4 |
| **Consciousness threshold (seen and correct)** | | | |
| **Group** | $F_{3,47} = 7.61$ | $P < 0.001$ | – |
| **Schizophrenia vs. healthy controls** | $t_{21} = 3.65$ | $P = 0.002$ | – |
| **OCD poor insight vs. healthy controls** | $t_{10} = 2.27$ | $P = 0.048$ | – |
| **OCD good vs. poor insight** | $t_{9} = -3.15$ | $P = 0.012$ | – |
| **OCD good insight vs. healthy controls** | $t_{22} = -2.15$ | $P = 0.043$ | – |
| **OCD poor insight vs. schizophrenia** | $t_{18} = -0.64$ | $P = 0.53$ | 1/BF = 2.3 |
| **Inflection point** | | | |
| **Group** | $F_{3,47} = 5.70$ | $P = 0.002$ | – |

| Measures | *F*-value | *p*-value | Bayes Factor |
|---|---|---|---|
| **Schizophrenia vs. healthy controls** | **$t_{23}$ = 3.30** | ***P* = 0.003** | – |
| **OCD poor insight vs. healthy controls** | $t_{10}$ = 1.61 | *P* = 0.14 | BF = 1.51 |
| **OCD good insight vs. healthy controls** | $t_{23}$ = -1.81 | *P* = 0.08 | 1/BF = 1.25 |
| **OCD poor insight vs. schizophrenia** | $t_{15}$ = -0.66 | *P* = 0.52 | 1/BF = 2.3 |
| **OCD good vs. poor insight** | **$t_{9}$ = -2.39** | ***P* = 0.041** | – |
| **Lower asymptote** | | | |
| **Group** | **$F_{3,47}$ = 0.49** | ***P* = 0.69** | 1/BF = 6.0 |
| **Upper asymptote** | | | |
| **Group** | **$F_{3,47}$ = 5.33** | ***P* = 0.003** | – |
| **Schizophrenia vs. healthy controls** | **$t_{22}$ = -3.00** | ***P* = 0.006** | – |
| **OCD poor insight vs. healthy controls** | **$t_{12}$ = -2.89** | ***P* = 0.013** | – |
| **OCD good insight vs. healthy controls** | $t_{21}$ = 0.46 | *P* = 0.65 | 1/BF = 2.5 |
| **OCD poor insight vs. schizophrenia** | $t_{23}$ = 0.41 | *P* = 0.69 | 1/BF = 2.6 |
| **OCD good vs. poor insight** | **$t_{11}$ = 3.23** | ***P* = 0.008** | – |
| **Slope** | | | |
| **Group** | **$F_{3,47}$ = 3.87** | ***P* = 0.015** | – |
| **Schizophrenia vs. healthy controls** | $t_{32}$ = 1.64 | *P* = 0.11 | 1/BF = 1.1 |
| **OCD poor insight vs. healthy controls** | **$t_{21}$ = 2.21** | ***P* = 0.038** | – |
| **OCD good insight vs. healthy controls** | $t_{22}$ = -1.50 | *P* = 0.15 | 1/BF = 1.5 |
| **OCD poor insight vs. schizophrenia** | $t_{21}$ = -0.64 | *P* = 0.53 | 1/BF = 2.4 |
| **OCD good vs. poor insight** | **$t_{15}$ = -3.89** | ***P* = 0.002** | – |
| **$R^2$** | | | |
| **Group** | **$F_{3,47}$ = 7.87** | ***P* < 0.001** | – |
| **Schizophrenia vs. healthy controls** | **$t_{29}$ = -2.98** | ***P* = 0.006** | – |
| **OCD poor insight vs. healthy controls** | **$t_{17}$ = -3.51** | ***P* = 0.003** | – |
| **OCD good insight vs. healthy controls** | $t_{17}$ = 1.51 | *P* = 0.15 | BF = 1.3 |

| Measures | *F*-value | *p*-value | Bayes Factor |
|---|---|---|---|
| **OCD poor insight vs. schizophrenia** | $t_{22}$ = 0.36 | *P* = 0.72 | 1/BF = 2.6 |
| **OCD good vs. poor insight** | **$t_{15}$ = 4.60** | ***P* < 0.001** | – |
| **Corrected consciousness threshold (seen and correct)** | | | |
| **Group** | **$F_{3,47}$ = 6.64** | ***P* < 0.001** | – |
| **Schizophrenia vs. healthy controls** | **$t_{23}$ = 3.37** | ***P* = 0.003** | – |
| **OCD poor insight vs. healthy controls** | $t_{10}$ = 1.94 | *P* = 0.081 | BF = 2.5 |
| **OCD good insight vs. healthy controls** | **$t_{22}$ = -2.10** | ***P* = 0.047** | – |
| **OCD poor insight vs. schizophrenia** | $t_{17}$ = -0.58 | *P* = 0.57 | 1/BF = 2.4 |
| **OCD good vs. poor insight** | **$t_9$ = -2.91** | ***P* = 0.017** | – |

### Supplementary Figures

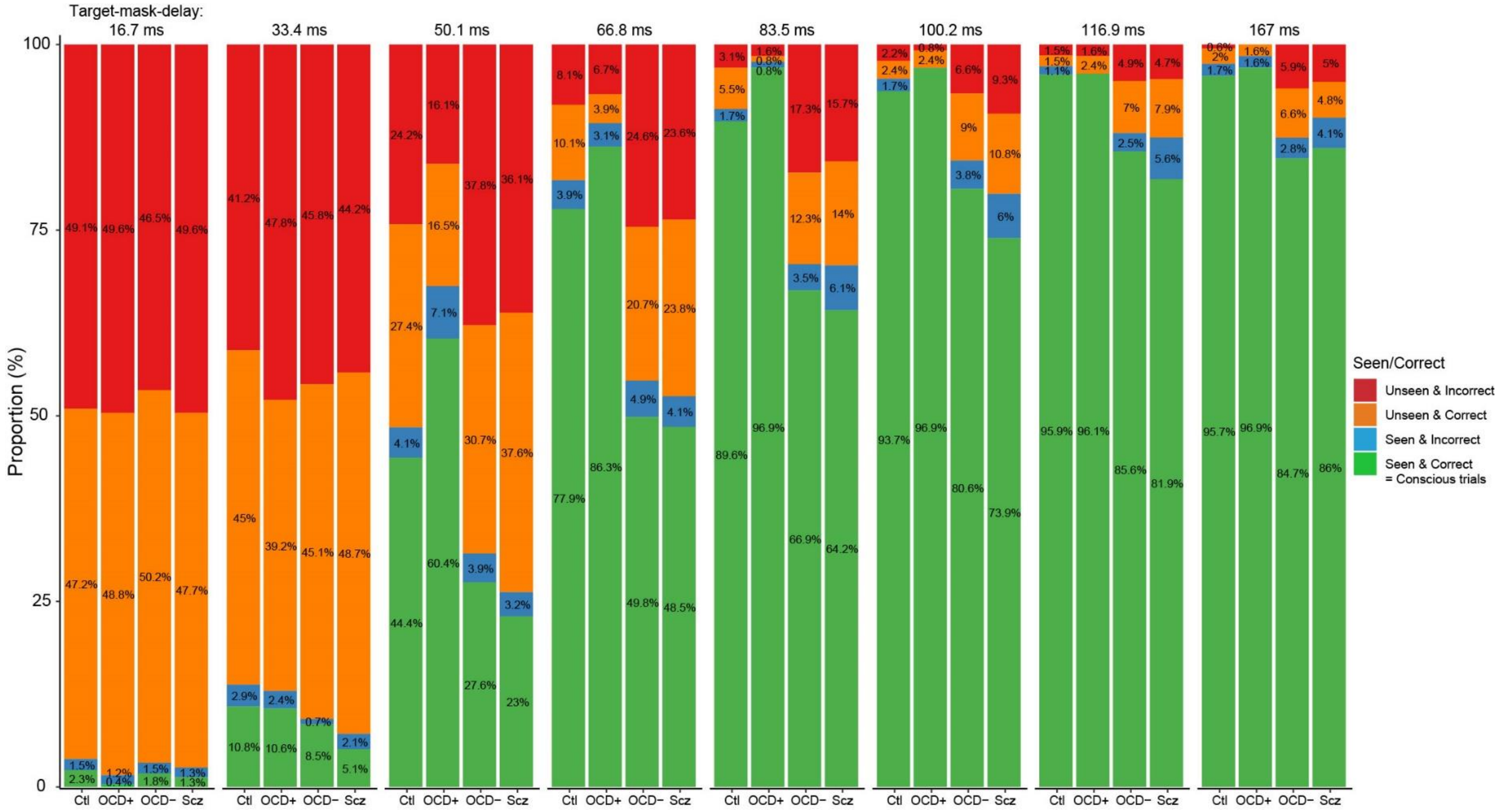


**Supplementary Fig. 1. Overview of participants' responses across target-mask delay** | Repartition of Seen × Correct responses by group and by target-mask delay (SOA). Proportion of Seen and Correct responses increase with target-mask delay. Patients with schizophrenia (Scz) and OCD with poor insight (OCD−) have significantly less "seen" and correct responses than healthy controls (Ctl) and patients with OCD with good insight (OCD+). Proportion of correct responses among unseen trials is about 50% in all groups, suggesting that differences in groups are not due to biases in response or different strategies of responses.

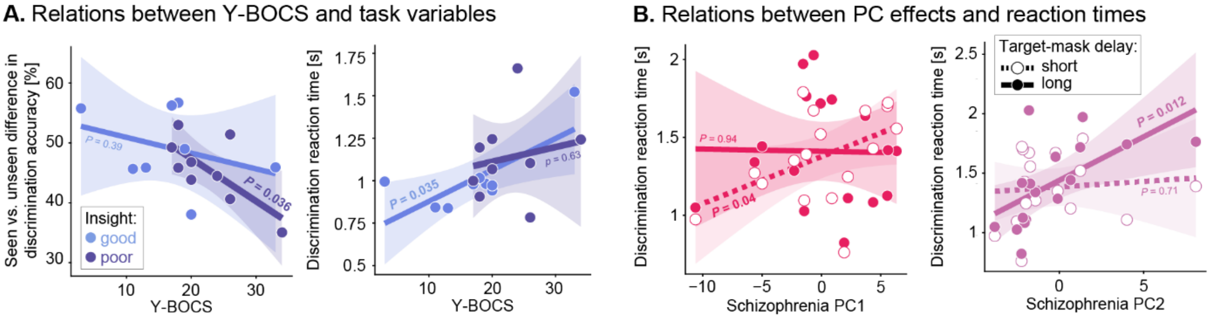


**Supplementary Fig. 2. Reaction times and metacognitive measures are modulated by Y-BOCS and schizophrenia PC** | Each dot represents a participant, lines represent linear regressions, and shaded areas represent 95% confidence intervals. **A**. Left. In the group of patients with OCD, Y-BOCS modulates seen vs. unseen difference in discrimination accuracy in patients with OCD and poor insight but not in patients with good insight. Right. By contrast, Y-BOCS modulates discrimination reaction times in patients with OCD and good insight but not in patients with poor insight. **B**. Left. In the group of patients with schizophrenia, PC1 modulates discrimination reaction times for short target-mask delay only. Right. By contrast, PC2 modulates discrimination reaction times for long target-mask delay only.
